# Immersive virtual prism adaptation therapy with depth-sensing camera: A feasibility study with functional near-infrared spectroscopy in healthy adults

**DOI:** 10.1101/2021.04.07.438873

**Authors:** Sungmin Cho, Won-Seok Kim, Jihong Park, Seung Hyun Lee, Jongseung Lee, Cheol E. Han, Nam-Jong Paik

## Abstract

Unilateral spatial neglect (USN) is common after stroke and associated with poor functional recovery. Prism adaptation (PA) is one of the most supported modality able to ameliorate USN but underapplied due to several issues. Using immersive virtual reality and depth-sensing camera, we developed the virtual prism adaptation therapy (VPAT) to overcome the limitations in conventional PA. In this study, we investigated whether VPAT can induce behavioral adaptations and which cortical area is most significantly activated. Fourteen healthy subjects participated in this study. The experiment consisted of four sequential phases (pre-VAPT, VPAT-10°, VPAT-20°, and post-VPAT) with functional near-infrared spectroscopy recordings. Each phase consisted of alternating target pointing and resting (or clicking) blocks. To find out the most significantly activated area during pointing in different phases (VPAT-10°, VPAT-20°, and Post-VPAT) in contrast to pointing during the pre-VPAT phase, we analyzed changes in oxyhemoglobin concentration during pointing. The pointing errors of the virtual hand deviated to the right-side during early pointing blocks in the VPAT-10° and VPAT-20° phases. There was a left-side deviation of the real hand to the target in the post-VPAT phase. The most significantly activated channels were all located in the right hemisphere, and possible corresponding cortical areas included the dorsolateral prefrontal cortex and frontal eye field. In conclusion, VPAT may induce behavioral adaptation with modulation of the dorsal attentional network. Future clinical trials using multiple sessions of a high degree of rightward deviation VPAT over a more extended period are required in stroke patients with unilateral spatial neglect.

## 1 Introduction

Stroke is the leading cause of acquired disability worldwide and causes various kinds of impairments (Feigin et al. 2014). Although motor impairment including hemiplegia is most common (Lawrence et al. 2001), recovery from non-motor impairments is also important and is targeted for rehabilitation in patients with stroke (Brewer et al. 2012). Unilateral spatial neglect (USN) (a deficit in attention to and awareness of the contralesional side of space), one such non-motor impairment, is common and persists usually after right hemisphere damage (Appelros et al. 2002; Buxbaum et al. 2004). Because USN is associated with poor functional recovery and lower social return, an appropriate rehabilitative intervention has to be timely provided (Jehkonen et al. 2000; Jehkonen, Laihosalo, and Kettunen 2006).

Among various rehabilitation approaches including both top-down approaches such as visual scanning and bottom-up approaches including prism adaptation (PA), caloric vestibular stimulation and neck vibration (Saevarsson, Halsband, and Kristjánsson 2011; Kerkhoff and Schenk 2012), PA is one of the most supported modality able to ameliorate USN (Yang et al. 2013), leading to recalibration of the egocentric coordinate system through repetitive pointing tasks to the deviated targets with prism glasses (Smit et al. 2013). The effects of PA on recovery from USN have been reported in previous studies of subacute or chronic stroke (Mizuno et al. 2011; Shiraishi et al. 2008). However, despite of the possible beneficial effect of PA on post-stroke USN, PA is underapplied in clinical practice (Maxton et al. 2013; Barrett, Goedert, and Basso 2012).

Despite its benefits, there are several issues regarding conventional PA therapy. In conventional PA therapy, patients wear prism glasses and perform repetitive pointing to targets deviated by the prism glasses while their hand trajectories to the targets should be masked before finally touching the targets. This conventional PA setting prevents its broad application. First, since the prism glasses have fixed parameters allowing for a certain visual displacement, PA requires multiple prism glasses, which is not practically and/or economically feasible. Although no dose-response studies have reported the effect of the degree of prismatic displacement on USN recovery, the degree of displacement has to be adjusted according to the patient’s ability to adapt, usually between 10° and 20° (Barrett, Goedert, and Basso 2012). In addition, minimizing self-corrections by gradually increasing the optical displacement during PA may induce larger after-effects (Michel et al. 2007). Second, the setting for masking the hand trajectory and visual targets is burdensome and requires a certain amount of space. This often impedes PA application to patients who cannot sit stably or control their head in a sitting position. Third, the exposure session during conventional PA has to be conducted under the supervision of a therapist, which limits its use in the ward or home. Fourth, PA’s degree of adaptation and after-effects are hard to quantify. If these problems can be solved, PA can become more beneficial to patients with extended PA sessions run in a more effective manner (minimizing self-corrections), resulting in long-lasting improvements (Newport and Schenk 2012).

We developed a new PA system, the “Virtual Prism Adaptation Therapy (VPAT),” using immersive virtual reality (VR) and a depth-sensing camera to overcome the limitations of conventional PA. VPAT deviates the hand trajectory during pointing through VR. Since VR can change the amount of visual displacement, and selectively show information, our VPAT does not require multiple glasses or a bulky system for masking the hand trajectory. This new system is easy to use, requires less supervisions, and automatically quantifies the degree of adaptations and after-effects. We designed an experiment with healthy individuals to investigate whether VPAT can induce similar effects compared to conventional PA. As a first surrogate outcome, the pointing errors from the target during the adaptation and post-adaptation period were employed. The second surrogate outcome was the cortical activation pattern during pointing with or without the prism mode, during the adaptation and post-adaptation periods. Functional near-infrared spectroscopy (fNIRS) was used to investigate possible neural substrates related to VPAT.

## 2 Materials and methods

### 2.1 Participants

Inclusion criteria for this study were healthy individuals aged between 18 and 50 years, right handed—assessed by the Edinburgh handedness inventory (Caplan and Mendoza 2011), feasibility to wear the Oculus Rift DK2 (Oculus VR, LLC, CA, USA), and the ability to detect the objects in the immersive VR. Subjects with any history of disease involving the central nervous system such as stroke, traumatic brain injury, or Parkinson’s disease were excluded. All subjects provided their written informed consent for participation. This study was carried out according to the Declaration of Helsinki and Good Clinical Practice Guidelines, and the protocol was approved by the Seoul National University Bundang Hospital Institutional Review Board. This study protocol and preliminary behavioral results without fNIRS data in four study subjects was published previously (Cho et al. 2020).

### 2.2 Concept of the VPAT system

VR systems with head mounted displays provide immersive visual feedback. The hand tracking with the head mounted display eliminates the gap between the virtual hand location and the location of the virtual object in virtual space. This clearly differ from the system using a 2D or 3D display, stylus, and mouse. Synchronization of the coordinate system of VR contents and hand tracking coordination can induce more realistic interactions with virtual objects. It is also possible to intentionally misalign the virtual hand and virtual object during the pointing action. This may be similar to a visual field shifting in the PA paradigm using prism goggles. However, while prism goggles shift all visual fields, our VPAT system only shifts the virtual hand trajectory in the immersive VR.

The virtual hand shift using our VPAT system induces initial pointing errors. In the condition without VPAT mode (no virtual hand shift), the subject directly points the virtual target (**Fig. 1a**). During the early VPAT period, the subject’s virtual hand is shifted to the right from the VR target (**Fig. 1b**), which might make the subject perceive the real target location to the left of the initial pointing location. During the late VPAT period, the subject adapts to VPAT mode and the pointing errors of the virtual hand to the target would diminish and finally disappear (**Fig. 1c**). After the VPAT mode is switched off, the subject’s real hand will point to the left of the target (**Fig. 1d**), if the adaptation effect of VPAT is similar to conventional PA therapy.

**Fig. 1.**
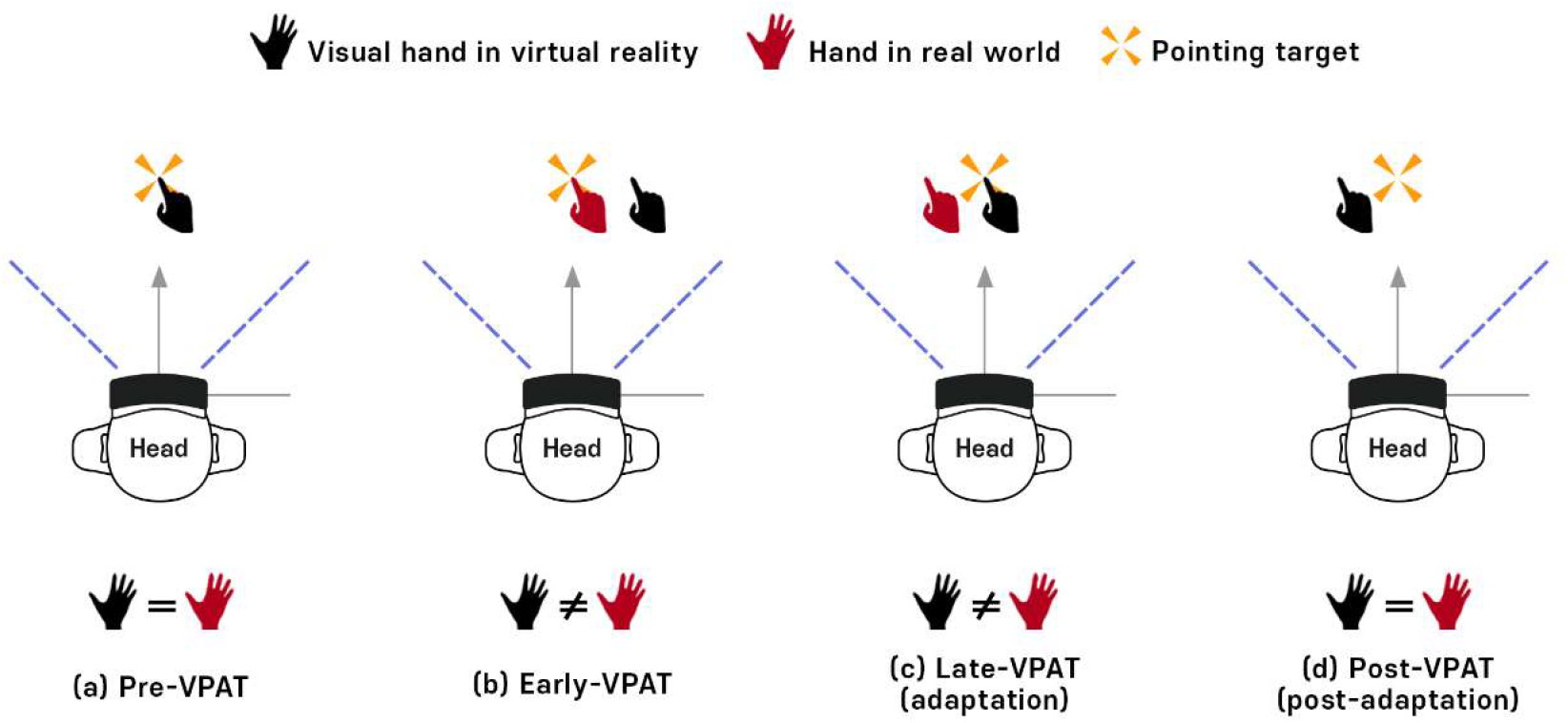
Concept of virtual prism adaptation therapy (VPAT) (a) Pre-VPAT: The position of the virtual hand in virtual reality and the real hand are always the same. (b) Early VPAT: Virtual hand shifted to the right side from the real hand. Virtual and real hand positions are not same. (c) Late VPAT (adaptation): After adaptation, the virtual hand will reach the target correctly, but the real hand will deviate to the left side from the object. (d) Post-VPAT (post-adaptation): Virtual and real hand are in the same position and the real hand still points the left direction from the target.

### 2.3 Hardware and software design of the VPAT system

The VPAT system uses the Oculus Rift DK2 as a head mounted display (HMD), while the Leap Motion sensor (Leap Motion, CA, USA) is mounted on the front of the HMD for hand tracking (**Fig. 2**). We configured a common desk environment with a virtual environment. Before projecting the user’s hands into the virtual space in real time, the hand skeleton dummies detected by the Leap Motion sensor are rotated based on the longitudinal axis for the visual hand shift.

**Fig. 2.**
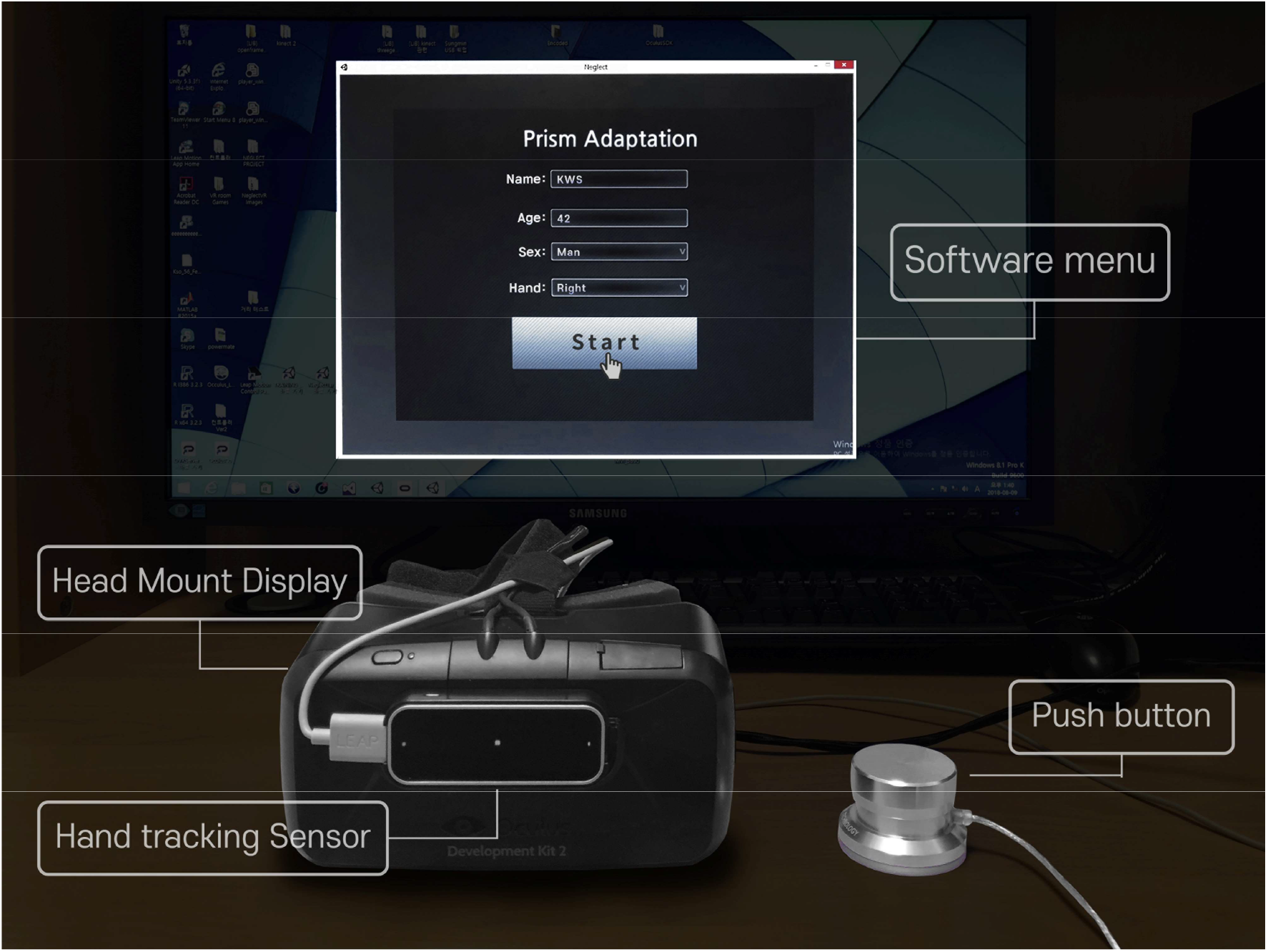
VPAT system configuration. **The** Occulus rift DK2 was used as a head mount display for VR rendering. Leap motion was used for hand tracking. A Push button was used during the clicking session in our experiment, and the software menu was constructed to save user’s information including name, age, gender, and the hand side used for training.

The location of a target object on a virtual table is defined as angle θ from the center of view. In the initial calibration phase, the distance of the object is specified as a comfortably reachable distance allowing the user to touch them (**Fig. 3a**). First, the coordinate systems between the head and hand should be co-aligned. The coordinate system of the visual hand is transformed by the angle φ in relation to the head center (**Fig. 3b**). If a point from the HMD coordinate system is ***p***_*x*|*hmd*_ and the point in the hand sensor coordinate system is ***p***_*x*|*hand*_, the relationship is given by the following equation:

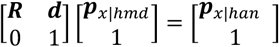

where **R** is a rotation matrix, and **d** the translation vector between the HMD coordinates and hand coordinates. Since the hand sensor is attached to the front of the HMD without rotation, **R** is the identity matrix, and **d** the relative position vector of the hand sensor from the HMD. Second, the experimental condition is given by ***R***_*shift*_ and ***d***_*shift*_. The hand shift was applied using the following equation:

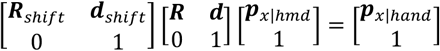

In our experiment, the hand only rotated horizontally. So, **R**_*shift*_ is

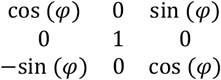

and ***d***_*shift*_ = **0**.

**Fig. 3.**
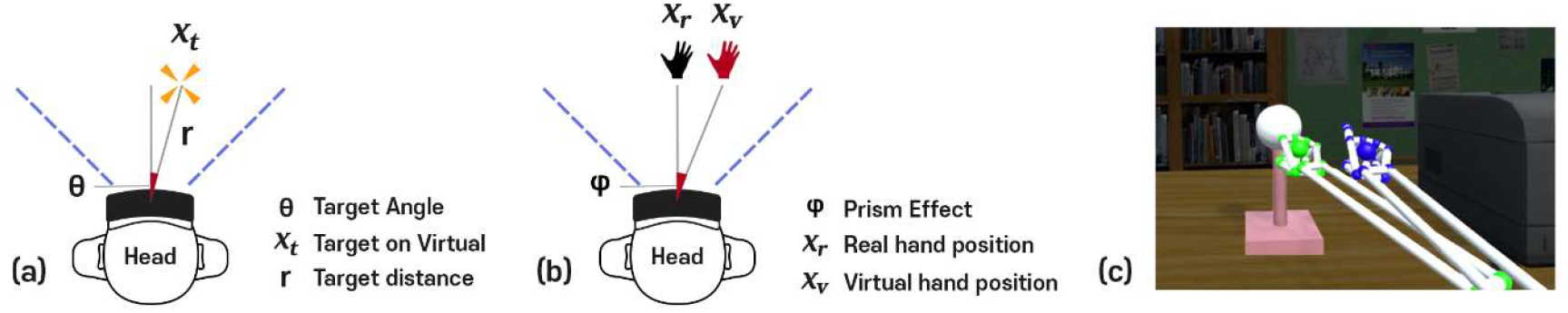
Hand trajectory deviations in VPAT system. (a) Target location (*x*_*t*_) defined as angle (*θ*) and distance (*r*) from initial calibration, (b) Virtual hand position (*x*_*v*_) shifted to the right side at a deviation angle(*φ*) from the real hand position (*x*_*r*_), (c)Hand trajectory: The green hand indicates the real hand trajectory and the blue hand indicates the virtual hand trajectory shifted to the right side.

### 2.4 Link between the VPAT system and fNIRS

The VPAT system was connected to the fNIRS system, NIRScout (NIRx Medical Technologies, LLC, MN, USA). The VPAT signal acted as a trigger and the events ecorded in the fNIRS system. The VPAT system was implemented with Unity 3D. The remote keyboard control software using TCP/IP communication was also implemented to synchronize the starting event with the fNIRS system. The fNIRS system is controlled by Superlab 5.0 (Cedrus Corporation, San Pedro, USA) as an event trigger method. The remote command key in a computer where Superlab 5.0 is installed was used to initiate the fNIRS recording.

### 2.5 Experimental design

Our protocol of video-recorded experiments was extensively documented previously (Cho et al. 2020). Briefly, subjects sat comfortably and wore the VPAT system. They pointed the target in the VPAT system without the prism mode for simple familiarization where they experience target shifts in the prism mode. Then the fNIRS cap with optodes in the montage set for this study was added. The experimental design is shown in **Fig. 4**. The experiments started from Phase 1, no VPAT mode. Four pointing and four clicking blocks appeared alternatively for 4 minutes. Phase 2 and Phase 3 were conducted with VPAT mode (Phase 2: 10° deviation, Phase 3: 20° deviation); each phase was composed of five pointing blocks and five resting blocks which appeared alternatively for total 5 minutes. Phase 4 was conducted without VPAT mode and represents the post-prismatic adaptation period. Phase 4 consists of five pointing and clicking blocks alternatively appearing for a total 5 minutes. In the clicking block, subjects were instructed to click the button as soon as possible when the visual target appeared at 3 second intervals, using their right index finger. Each clicking block consisted of a total of 10 clicks for 30 seconds. The pointing task block was composed of 10 pointings for 30 seconds in total. Subjects were instructed to point the visual targets with the right index finger as fast as possible. The visual target was presented on a position deviated 10° -right or 10°-left from the midline at 3 second intervals and in random orders. When the subject correctly pointed the target, the color changed to red.

**Fig. 4.**
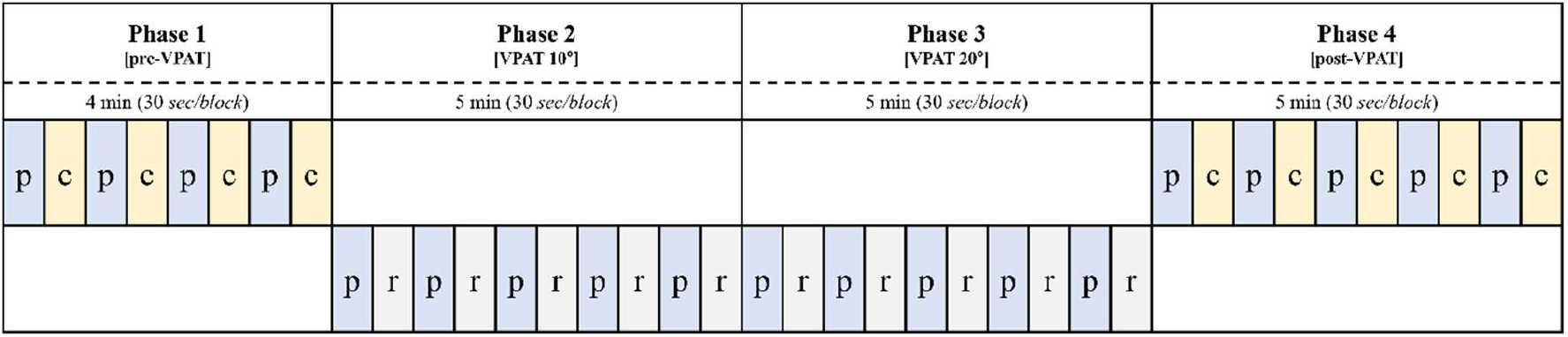
Experimental design. VPAT, virtual prism adaptation therapy; P pointing; C, clicking; R, resting.

### 2.6 fNIRS measurement

A continuous-wave type fNIRS system was used to measure changes in oxygenated hemoglobin [HbO] caused by cortical activations during the experiment. The manufacturer’s NIRStar 14.1 program was operated to control the NIRScout. The fNIRS system uses two wavelengths of near-infrared light, 760 and 850 nm, for measurement and records raw optical density data with a sampling rate of 4.17 Hz. A total of 39 measurement channels were formed by positioning 15 source and 13 detector LED optodes (optical probes). The optode positions were designed to mainly cover the dorsal frontoparietal network and the motor cortices of both hemispheres (**Fig. 5**), based on the international 10-20 system (Klem et al. 1999). The separation between each source and detector was approximately 3 cm, enabling the near-infrared light to reach cortical areas. For optode fixation, textile EEG caps (EASYCAP, Herrsching, Germany) in three different sizes (54, 56, and 58 cm circumference) were prepared and selected depending on subject’s head size.

**Fig. 5.**
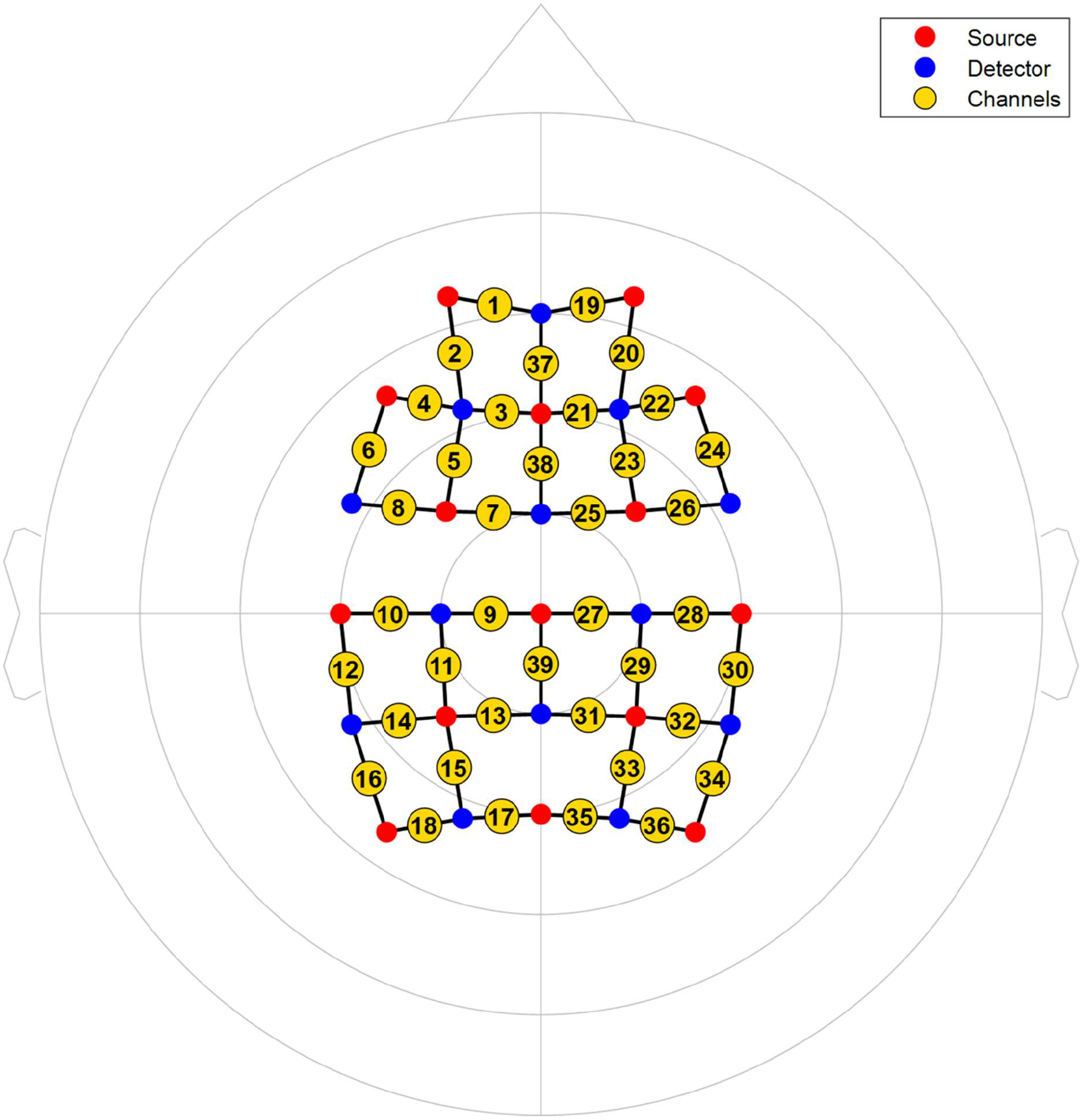
fNIRS recording montages for the experiment. Red circles indicate light sources, blue circles detectors, and yellow circles channels.

### 2.7 Pointing error analysis

All hand trajectory data was recorded with a sampling rate of 60 Hz during the experiment. After the target appears, pointing errors were calculated using the target angle and the detection of the moment at which the hand appears, with the position of the head as the reference point. The first pointing errors during each block in the four phases were used for analysis. They are presented using a box and whisker plot. Multiple Wilcoxon signed-rank tests were used to test the differences between six time points (three transitions between phases and three time points between the first and last block in phases 2 to 4). A 2-tailed Bonferroni correction, with a *p* < 0.008 considered statistically significant, was performed to adjust the type I error due to multiple comparisons. Statistical analyses were performed using the PASW statistical package (SPSS version 18.0, SPSS).

### 2.8 fNIRS data processing and analysis

Preprocessing was performed using the nirsLAB program, version 2019.04 (NIRx Medical Technologies LLC, MN, USA). The optical density raw data was converted to [HbO] by applying the modified Beer–Lambert Law (Delpy et al. 1988). Then, the resultant [HbO] was bandpass filtered with a passband between 0.009 and 0.2 Hz to remove drifts and task-unrelated physiological effects, such as respiration and cardiac activities (Tachtsidis and Scholkmann 2016). [HbO] levels were only used for analysis because it is more sensitive to the cortical blood flow changes according to the task and deoxyhemoglobin has shown considerable individual differences in task-related changes (Strangman, Franceschini, and Boas 2003).

General linear model (GLM)-based statistical data analysis for the task “pointing’” was performed using Statistical Parametric Mapping (SPM, version 8) (Penny et al. 2011), integrated within the nirsLAB program. In Level-1 analysis (within-subject), the GLM coefficients of [HbO] in each channel were estimated with the canonical hemodynamic response function (HRF) after temporal filtering by discrete cosine transform function and, in turn, precoloring by HRF was implemented. Then, the three t-contrasts were specified for the comparisons of the pointing blocks in each of the three phases (VPAT-10°, VPAT-20°, Post-VPAT) against the pointing blocks in the default phase, pre-VPAT. Subsequently, for each subject, SPM t-maps were computed based on those t-contrasts with p<0.05 (uncorrected). In Level-2 analysis (group), group t-maps for the three comparisons were generated to identify the channels most significantly activated by pointing, in contrast to pre-VPAT pointing (VPAT-10° vs. Pre-VPAT, VPAT-20° vs. Pre-VPAT, Post-VPAT vs. Pre-VPAT).

## 3 Results

### 3.1 Pointing errors during the experiment

Fourteen healthy subjects (seven male, seven female) participated in this study. Their ages were 27.8±4.1 years (range: 22 – 39). Pointing errors in each block were presented using a box and whisker plot (**Fig. 6**). Pointing errors were around 0° in the pre-VPAT phase. In the VPAT-10° phase, the errors initially deviated towards the right with a median value of 6.32° and interquartile range (IQR) of 3.00°and gradually decreased to 2.41° (IQR: 2.33°). At the start of the VPAT-20° phase, pointing errors moved rightward 8.10° (IQR: 3.00°) and decreased to 4.43° (IQR: 4.23°). In the post-VPAT phase, there was a substantial leftward pointing error (8.48°, IQR: 5.69°) in the first block of pointing, decreasing gradually afterward. The differences in pointing errors between the three phase transitions were statistically significant (*p*<0.008), as were the changes between the first and last block in the three phases (VPAT-10°, 20°and post-VPAT phase) (*p*<0.008) (**Fig 6**).

**Fig. 6.**
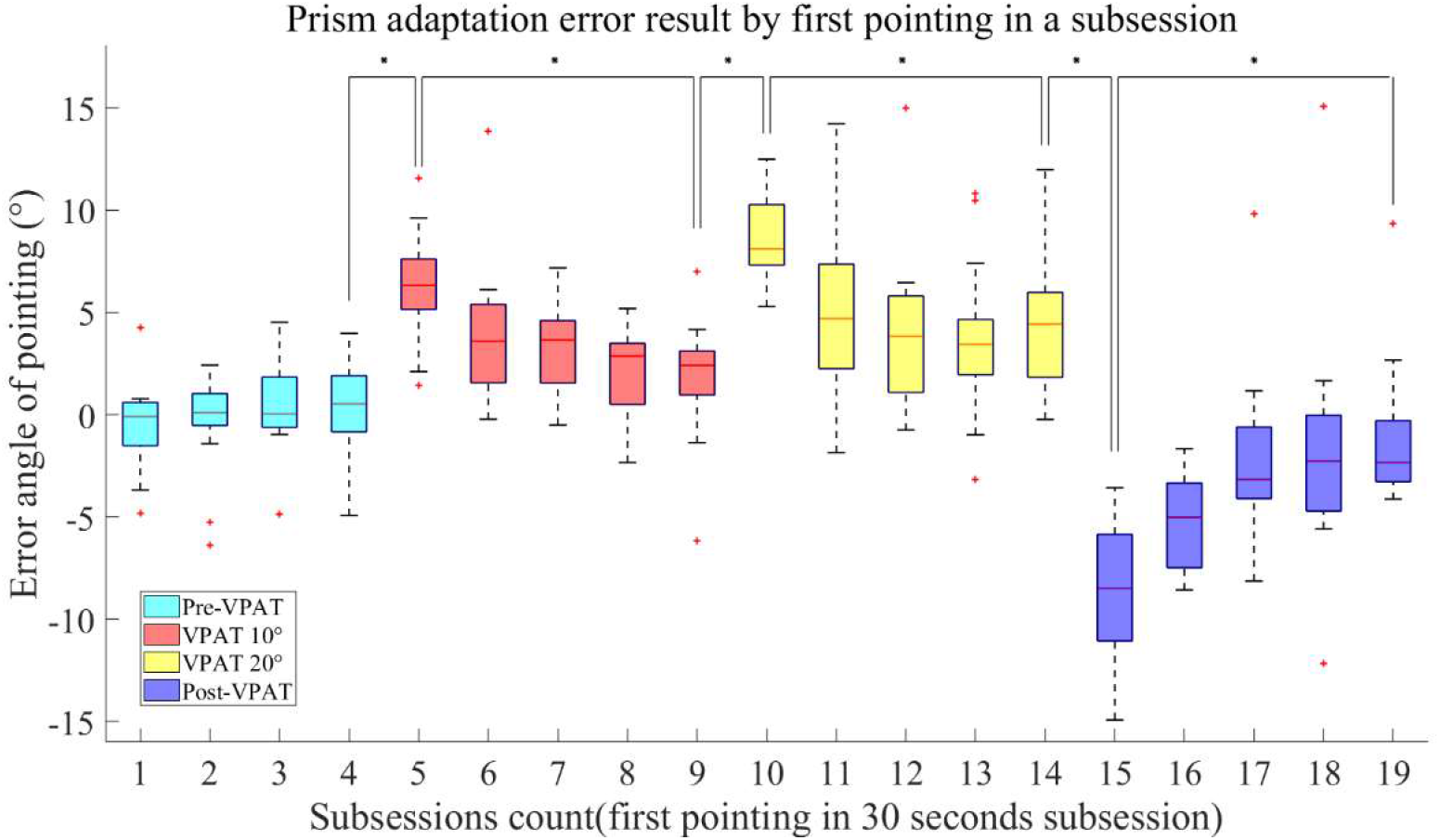
Pointing errors during the experiment. Positive value indicates rightward deviation, whereas negative value indicates leftward deviation. VPAT, virtual prism adaptation therapy. ^*^*p*<0.008 by Wilcoxon signed rank test with Bonferroni correction.

### 3.2 Cortical activation as measured by fNIRS

The cortical areas most significantly activated during pointing were identified by using contrast to pointing trials during pre-VPAT in the following three phases: VPAT-10° (channel 20, *p*=0.108), VPAT-20° (channel 24, *p*=0.027) and post-VPAT (channel 22, *p*=0.221) (**Table 1**). The most significantly activated channels were all located in the right hemisphere, and possible corresponding cortical areas include the dorsolateral prefrontal cortex and the frontal eye field (Brodmann areas 8, 9 and 46) (Zimeo Morais, Balardin, and Sato 2018).

**Table 1.** Cortical regions most significantly activated during virtual prism adaptation therapy (VPAT).

| Condition | Side | Channel no. | t | p |
| --- | --- | --- | --- | --- |
| (VPAT-10° pointing) – (Pre-VPAT pointing) | Right | 20 | 1.663 | 0.108 |
| (VPAT-20° pointing) – (Pre-VPAT pointing) | Right | 24 | 2.342 | 0.027 |
| (Post-VPAT pointing) – (Pre-VPAT pointing) | Right | 22 | 1.254 | 0.221 |

## 4 Discussion

The mechanisms used in the VPAT system differ from real prism therapy. In real PA, the whole visual field is shifted to the right. However, in our VPAT system, only the virtual hand trajectory is deviated to the right, while the actual target remains in its real position (**Fig. 3**) (Aasted et al. 2015). During VPAT mode, the behavioral adaptation during pointing was observed according to the degree of virtual hand deviation (**Fig 6**). In recent studies with a VPAT system using HMD and a hand-held controller, healthy adults or stroke patients with USN showed clear post-VPAT adaptations in open-loop pointing (Gammeri et al. 2020; Bourgeois et al. 2021). The pointing errors in post-VPAT adaptation at 20° deviation was about 8° in healthy adults (Gammeri et al. 2020), and that after 15° deviation VPAT was 4.4° in stroke patients with USN (Bourgeois et al. 2021), similar to our study (8.48°). Although the transfer effect of VPAT was not investigated in our study, Gammeri et al. showed that the pointing bias after VPAT was a significant predictor of transfer to the bisection task in healthy adults.

Two previous studies evaluated the transfer effect of VPAT using bisection, landmark or cancellation tasks but their results were contradictory (Gammeri et al. 2020; Bourgeois et al. 2021). The transfer effect was observed after 30° VPAT adaptation in healthy adults (Gammeri et al. 2020), but not in the cross-over study among stroke patients with USN (Bourgeois et al. 2021). We think that the negative results of VPAT in the study by Bourgeois et al. (Bourgeois et al. 2021) with stroke patients should be cautiously interpreted. First, the included stroke patients were heterogenous in terms of the USN phenotype. They conducted screening using the confrontation test or perimetry, and eight among 15 patients showed hemianopsia, which might not be differentiated from pure USN. Prism are usually employed for partially extending the visual field in hemianopsia (Grunda, Marsalek, and Sykorova 2013) but PA to deviate the visual field has no sound rationale for the recovery of hemianopsia after stroke, therefore, possibly lessening the effect of VPAT. In addition, patients with both allocentric and/or egocentric USN were included. In some patients, only the line bisection test showed pathological performances while cancellation tests were normal, which translates as allocentric USN rather than egocentric USN. Considering the possible different effects of PA between egocentric and allocentric neglect (Mancuso et al. 2018; Gossmann et al. 2013), the heterogenous USN phenotypes in Bourgeois et al.’s study (Bourgeois et al. 2021) might dampen VPAT’s effects. Furthermore, PA can be more beneficial in patients with motor-intentional aiming deficits than in those with predominantly perceptual USN (Goedert et al. 2014), which were not considered in the previous study (Bourgeois et al. 2021). Second, PA may act on USN due to cumulative effects. For instance, Nys et al. showed that the performance before and after the PA session on each day did not change, despite a significant improvement in spatial tasks after four daily consecutive PA sessions (Nys et al. 2008). In addition, the transfer effect of PA may be larger 2 hours after PA than immediately after (Rossetti et al. 1998), thus the lack of significant transfer effect immediately after one VPAT session is not sufficient to conclude that VPAT is not effective on USN. Finally, the effect of PA can be influenced by damaged brain areas due to stroke (Gutierrez-Herrera et al. 2020). Therefore, further clinical trials to investigate the VPAT’s efficacy on USN are required. Specifically, a future trial should consider conducting multiple VPAT sessions over an extended period to prove the long-term cumulative effects (Frassinetti et al. 2002; Serino et al. 2007) and offer more intensity in terms of the degree of rightward deviations, considering the threshold of deviations for the immediate VPAT aftereffects (Gammeri et al. 2020). In addition, a multicenter trial may have to be considered, due to the relatively small effect sizes of PA (Umeonwuka, Roos, and Ntsiea 2020) and higher severity and older ages in stroke patients with USN, preventing them to participate PA trials (Gottesman et al. 2008).

Our second purpose was to investigate which is the most relevant cortical area related to VPAT. Since the induction mechanism of behavioral adaptations in our VPAT was different from the conventional therapy using a real prism, we did not know if the activated cortical areas would be similar to previous studies with a real prism, due to lack of studies on this aspect. In our study, the most activated areas during and after VPAT included the right frontal eye field and the dorsolateral prefrontal cortex (**Table 1**) (Zimeo Morais, Balardin, and Sato 2018), although without statistical significance in the VPAT-10° and post-VPAT periods. The frontal eye field is an area involved in the dorsal attentional network, which mediates top-down stimulus-response selection (Corbetta et al. 2005), producing inattention to an object in the contralesional far-space when that area is damaged (Rizzolatti, Berti, and Gallese 2000). A recent functional MRI study found that rightward PA increased resting functional connectivity in healthy adults between the right frontal eye field and right anterior cingulate cortex after PA, which may mean more attention to the left visual field (Tsujimoto et al. 2019).

In our study, the activated area was the right dorsolateral prefrontal cortex, not the left. A previous functional MRI study in patients with right hemispheric damage showed that rightward PA increased activation in the left prefrontal cortex (Crottaz-Herbette et al. 2017). It was postulated that PA induced bottom-up activation of the prefrontal cortex, which then led to the enhanced function of the left hemisphere to perceive the left visual field (Panico, Rossetti, and Trojano 2020). However, the role of the left prefrontal cortex on PA’s effect have not been well elucidated. Bottom-up stimulus driven control is known to be related with the ventral attentional network usually including the ventral prefrontal cortex, not the dorsal one (Corbetta and Shulman 2002). In addition, despite an ongoing debates, the dorsolateral prefrontal cortex is considered to be included in the dorsal attentional network (Corbetta and Shulman 2002; Qian et al. 2020). The shift from using the left ventral attentional network by PA, secondarily activates the right dorsal attentional network to perceive the left visual field, thus in our study, VPAT may induce the activation of the dorsal attentional network including both the right dorsolateral prefrontal cortex and frontal eye field. The various subject characteristics (right prefrontal cortex damaged patients vs. healthy adults) could also result in the different activation side in the prefrontal cortex between our study and that by Croattaz-Herbette et al. (Crottaz-Herbette et al. 2017). Due to these uncertainties in the role of the prefrontal cortex on PA and the difficulties in directly measuring the hemodynamic changes of the deeply located ventral prefrontal cortex by fNIRS, further research is warranted.

This study has several limitations. First, the exact localization of involved cortical areas using fNIRS was difficult without co-registration of each channel with an individual magnetic resonance image, even though we followed the 10-20 EEG channel position using an appropriately sized cap according to the subject’s head size. In addition, deeply located structures possibly related with the effect of PA such as the anterior cingulate cortex (Danckert, Ferber, and Goodale 2008) could not be directly monitored by fNIRS. Second, the single statistically activated channel was only found in the VPAT-20° condition, although most significantly activated channels were consistently located around the right dorsolateral prefrontal cortex and frontal eye field. This can be explained by PA’s dose-response effect. Gammeri et al. reported that only VPAT-30° could significantly induce the transfer effect but not VPAT-10° and VPAT-20° (Gammeri et al. 2020; Bourgeois et al. 2021). In addition, a total of 50 pointing with 30 sec resting time after each 10 pointings were used in our experiment, which was lower than the number of consecutive pointing (one hundred) in previous VPAT studies (Gammeri et al. 2020; Bourgeois et al. 2021). Finally, in situations where the hand was not detected properly due to occlusions or self-occlusions, there were often cases where the hand did not appear, even though it actually touched the virtual object during VPAT. Using VR controllers would reduce this error. However, fatigue might occur due to the holding pose and the controller weight. Hand tracking technology using multiple cameras on the front of the HMD could reduce errors caused by occlusion without fatigue (Han et al. 2020).

## Conclusions

VPAT deviating the hand trajectory rightward induces visuomotor adaptations. The most activated cortical areas during and immediately after VPAT were the dorsolateral prefrontal cortex and the frontal eye field, which are associated with the dorsal attentional network. The VPAT system has the following advantages compared to conventional PA: 1) easier adjustment of deviation angles, 2) gradual adjustment of deviations according to the patient’s adaptation, 3) blockage of external visual cues, and 4) quantification and monitoring of pointing errors during therapy. In times of COVID-19, the VPAT system can help patients and therapists set up the treatment environments at the patient’s home, with monitoring or self-guided algorithms (e.g., automatically adjusting the deviations according to the patient’s behavioral adaptation). Therefore, a future clinical trial using VPAT with a high degree of rightward deviation and multiple sessions over a more extended period is required in stroke patients with USN.

## Acknowledgements

Not applicable.

## Author Contributions

NJP and WSK supervised this study. WSK, NJP, and SC conceptualized and designed the study. WSK acquired the funding and provided the resources for the study. SC developed the VPAT system and WSK and JP suggested the rehabilitation contents in the system. JP, SHL, and JL supervised fNIRS data acquisition. JL processed fNIRS data and analyzed them. SC extracted the pointing error data and analyzed them. SC, WSK, JP, SHL, CEH, JL, and NJP interpreted the results from the data. WSK, SC, JP, JL, and CEH drafted the original manuscript. All authors read and revised the manuscript critically for important intellectual content and approved the final manuscript for publication. All authors agree to be accountable for all aspects of the work.

## Data Availability

The datasets generated and analyzed during the current study are available from the corresponding author on reasonable request.

## Declarations

### Ethics approval and consent to participate

Written, informed consent was obtained from each subject. The Seoul National University Bundang Hospital Institutional Review Board (IRB) approved this study protocol (IRB number: B-1605/345-006).

### Consent for publication

Not applicable.

